# CTCF is essential for proper mitotic spindle structure and anaphase segregation

**DOI:** 10.1101/2023.01.09.523293

**Authors:** Katherine Chiu, Yasmin Berrada, Nebiyat Eskndir, Dasol Song, Claire Fong, Sarah Naughton, Tina Chen, Savanna Moy, Sarah Gyurmey, Liam James, Chimere Ezeiruaku, Caroline Capistran, Daniel Lowey, Vedang Diwanji, Samantha Peterson, Harshini Parakh, Ayanna R. Burgess, Cassandra Probert, Annie Zhu, Bryn Anderson, Nehora Levi, Gabi Gerlitz, Mary C. Packard, Katherine A. Dorfman, Michael Seifu Bahiru, Andrew D. Stephens

**Affiliations:** Biology Department, University of Massachusetts Amherst, Amherst, MA 01003, USA; Biology Department of Molecular Biology, Faculty of Life Sciences, Ariel University, Ariel 40700, Israel; Molecular and Cellular Biology, University of Massachusetts Amherst, Amherst, MA 01003, USA

**Keywords:** CTCF, mitosis, DNA/chromatin, mitotic spindle, nucleus

## Abstract

Mitosis is an essential process in which the duplicated genome is segregated equally into two daughter cells. CTCF has been reported to be present in mitosis but its importance for mitotic fidelity remains to be determined. To evaluate the importance of CTCF in mitosis, we tracked mitotic behaviors in wild type and two different CTCF CRISPR-based genetic knockdowns. We find that knockdown of CTCF results in prolonged mitoses and failed anaphase segregation via time lapse imaging of SiR-DNA. CTCF knockdown did not alter cell cycling or the mitotic checkpoint, which was activated upon nocodazole treatment. Immunofluorescence imaging of the mitotic spindle in CTCF knockdowns revealed disorganization via tri/tetrapolar spindles and chromosomes behind the spindle pole. Imaging of interphase nuclei showed that nuclear size increased drastically, consistent with failure to divide the duplicated genome in anaphase. Population measurements of nuclear shape in CTCF knockdowns do not display decreased circularity or increased nuclear blebbing relative to wild type. However, failed mitoses do display abnormal nuclear morphologies relative to successful mitoses, suggesting population images do not capture individual behaviors. Thus, CTCF is important for both proper metaphase organization and anaphase segregation which impacts the size and shape of the interphase nucleus.

## Introduction

Mitosis is the process by which the duplicated genome is segregated between two daughter cells. This process requires the proper compaction of chromatin into chromosomes, alignment of chromosomes by the mitotic spindle in metaphase, and segregation of the chromosomes in anaphase. Most forms of life rely on mitosis to grow and replenish lost cells. Failure of mitosis can cause alterations in copy number of genes that can promote human diseases like cancer (Levine and Holland 2018). This can occur on a spectrum between loss of a single chromosome to complete failure of mitosis resulting in a tetraploid cell with 4 copies of the genome. Even though we have studied this phenomenon for decades, the list of proteins contributing to a successful mitosis continues to increase.

Many chromatin proteins have a role in interphase chromatin organization and expression while having a separate role during mitosis. An example is cohesin which provides chromatin looping in interphase to organize the genome (Seitan et al. 2013; Banigan et al. 2022) and holds duplicated sister chromatids together during metaphase (Guacci et al. 1997; Michaelis et al. 1997). We hypothesized that CTCF, another chromatin protein that has a major role in chromatin looping and interphase genome organization (Hansen et al. 2017), could have a role in mitosis as well. CTCF has been reported to continue to bind to chromatin throughout mitosis (Burke et al. 2005). One study reports that CTCF is important for recruiting CENP-E to the pericentromere in mitosis (Xiao et al. 2015) where CTCF is reportedly located via ChIP and immunofluorescence (Rubio et al. 2008). Interestingly, other studies have reported CTCF localized to centrosomes and/or the midzone (Zhang et al. 2004) and interacts with kaiso which also localizes to pericentriole (Defossez et al. 2005). Thus, while CTCF is present on mitotic chromosomes and in the mitotic spindle, whether it contributes to mitosis remains unclear.

To determine if CTCF has a role in mitotic fidelity, we used CTCF CRSIPR-based knockdown (KD) clones in B16 mouse melanoma cell line (Kaczmarczyk et al. 2022). We first live-cell imaged mitotic behavior by tracking SiR-DNA labeled in wild type and two different CTCF knockdowns. Loss of CTCF results in metaphase failures, anaphase segregation failures, and tripolar spindles. However, the spindle checkpoint and mitotic cycling remain intact. Population images of the mitotic spindle confirm that both gross chromosome alignment and miotic spindle structure were perturbed in a subset of CTCF knockdowns. Imaging of interphase nuclei further revealed that CTCF knockdown displayed abnormal nuclear size and shape post mitotic failure. Thus, CTCF is important for both proper metaphase organization and anaphase segregation which impacts the resulting interphase nucleus.

## Results

### CTCF is required for proper mitotic fidelity via metaphase alignment and anaphase segregation

Identifying the importance of CTCF in mitosis requires a CTCF knockdown (KD) cell line. Previously we generated CRISPR-based knockdown of CTCF in B16-F1 mouse melanoma cells to decrease the total CTCF protein levels by 60-70% while the remaining CTCF was truncated with reduced chromatin binding capabilities (Kaczmarczyk et al. 2022). Complete knockdown of CTCF was shown to be lethal (Wan et al. 2008; Moore et al. 2012; Nora et al. 2017), thus preventing the study of cell proliferation and mitosis. Therefore, the small amount of truncated CTCF in the clones we generated seems to enable their proliferation and studying mitosis. Specifically, CTCF knockdown clones were differently composed of truncations with a molecular weight of 120 and 65 KDa in clone c13 and 100 and 65 KDa in clone c21. Thus, these unique cell lines provide the ability to investigate CTCF knockdowns relative to their wild type parent cell line.

To determine if CTCF is essential for mitotic fidelity, we live-cell imaged wild type and CTCF knockdown cells over 24 hours using SiR-DNA (**Figure 1**). Wild type B16-F1 cells present low levels of abnormal mitotic behavior (5 ± 3%, **Figure 1B**). Both CTCF CRSIPR-based knockdown clones (c13 and c21) displayed a significant increase in failed mitoses (63 ± 9% and 31 ± 2% respectively, **Figure 1B**). Thus, our data reveal that CTCF has an essential role in mitotic fidelity and upon disruption results in mitotic failure.

**Figure 1.**
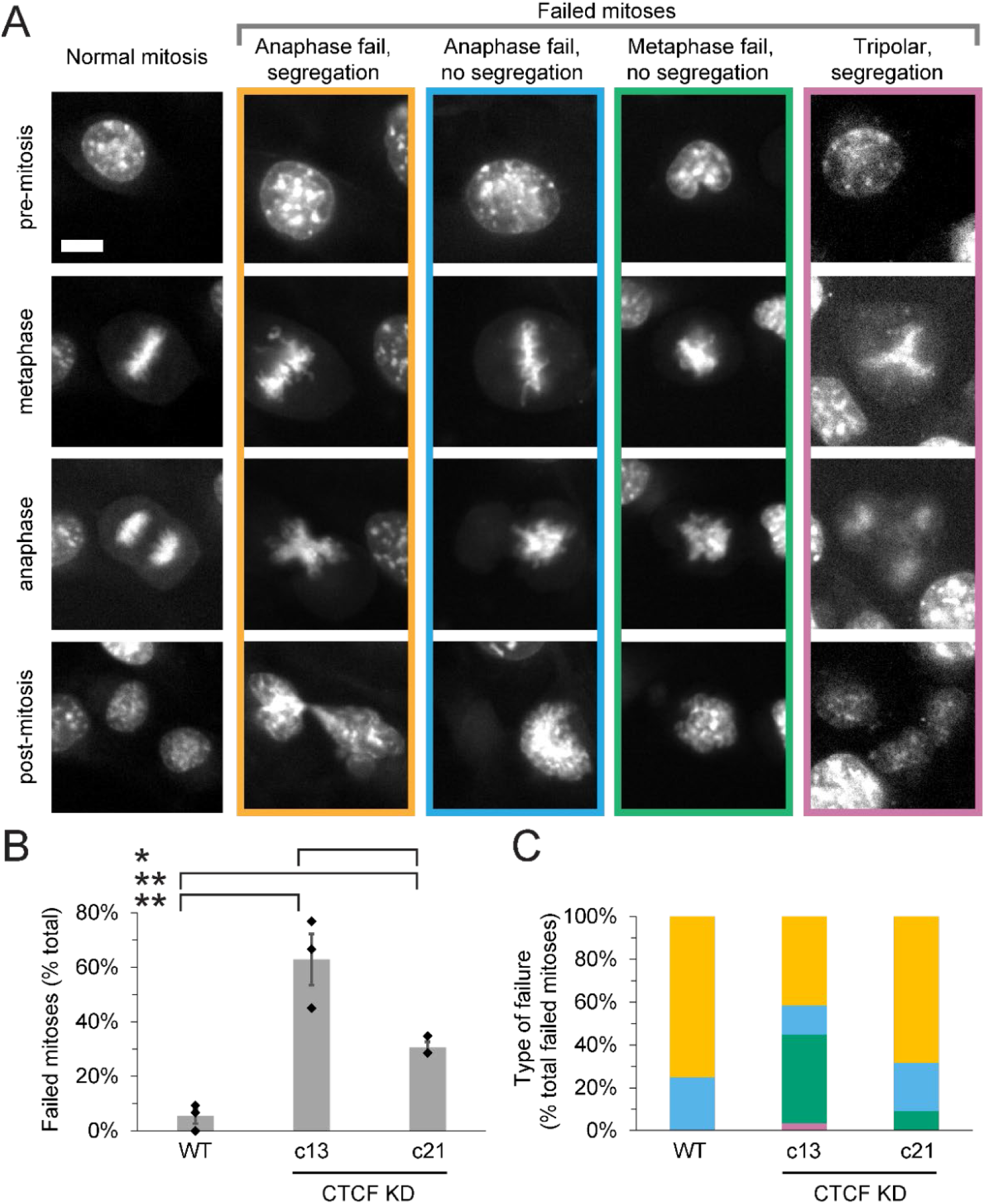
CTCF knockdowns display increased mitotic failure. (A) SiR-DNA labeled representative images of normal mitosis and different types of mitotic failures taken from overnight timelapses of B16 wild type (WT) and CTCF CRISPR knockdown clones (CTCF KD c13 and c21). See **Movies 1-5**. (B) Graphs of the percentage of mitotic failures and (C) the type of mitotic failure for wild type (WT) and CTCF CRISPR knockdown mutants (c13 and c21). Three biological replicates reported as mitotic failure out of all WT (1/15, 0/20, 3/32), CTCF knockdown c13 (10/13, 10/15, 9/20), and CTCF knockdown c21 (6/21, 8/28, 8/23). Error bar represents standard error. statistical tests are Student’s t-tests, significance denoted by * p < 0.05, ** p < 0.01, and *** p < 0.001, while no significance is denoted by ns. Scale bar is 10 µm.

Time lapse imaging allows us to determine when and how the mitotic failures occur during mitosis. We classified mitotic failure into four categories: anaphase failure with segregation, anaphase failure without segregation, metaphase failure without segregation, and tripolar segregation (**Figure 1, A and C; Movies 1-5**). Low levels of mitotic failure in wild type were due largely to anaphase bridges, where only one observed event showed failure to separate (1/71 mitotic events). Oppositely, the majority of events (55%) in CTCF knockdown c13 resulted in genome segregation failure in anaphase (**Figure 1C**). In anaphase-based segregation failures, cytokinesis was observed for all imaged events. The other CTCF knockdown clone c21, most of the events displayed some degree of segregation. Though cytokinesis appears to force segregation of the genome in many cases, and thus CTCF knockdowns present a cut phenotype (Samejima et al. 1993). Taken together, this data supports that CTCF is required for both successful metaphase alignment and anaphase segregation.

### Spindle checkpoint and mitotic cycling are maintained in CTCF knockdowns

Failure of the spindle assembly checkpoint (SAC) could be responsible for mitotic failure by not delaying in metaphase to correct erroneous attachments and alignment before proceeding into anaphase. To determine if the mitotic checkpoint is activated properly in CTCF mutants, we treated cells overnight with increasing concentrations of nocodazole to depolymerize microtubules that compose the mitotic spindle resulting in mitotic arrest (Harper 2005). Mitotic arrest was determined by joint imaging of SiR-DNA of condensed chromosomes and transmitted light showing a rounded mitotic cell (yellow arrows, **Figure 2A**). Wild type and CTCF knockdowns showed a moderate and similar percentage of cells arrested in mitosis upon low treatment of nocodazole (6 ± 1% 100ng/mL, **Figure 2, A and B**). Upon overnight treatment with 10-fold more nocodazole, wild type and both CTCF knockdowns showed a significant increase in mitotic arrest up from 6% to ∼21% (**Figure 2, A and B**). This data suggests that CTCF mutants have an intact spindle checkpoint. Thus, mitotic failure in CTCF knockdowns is likely not due to disruption of the mitotic spindle checkpoint.

**Figure 2.**
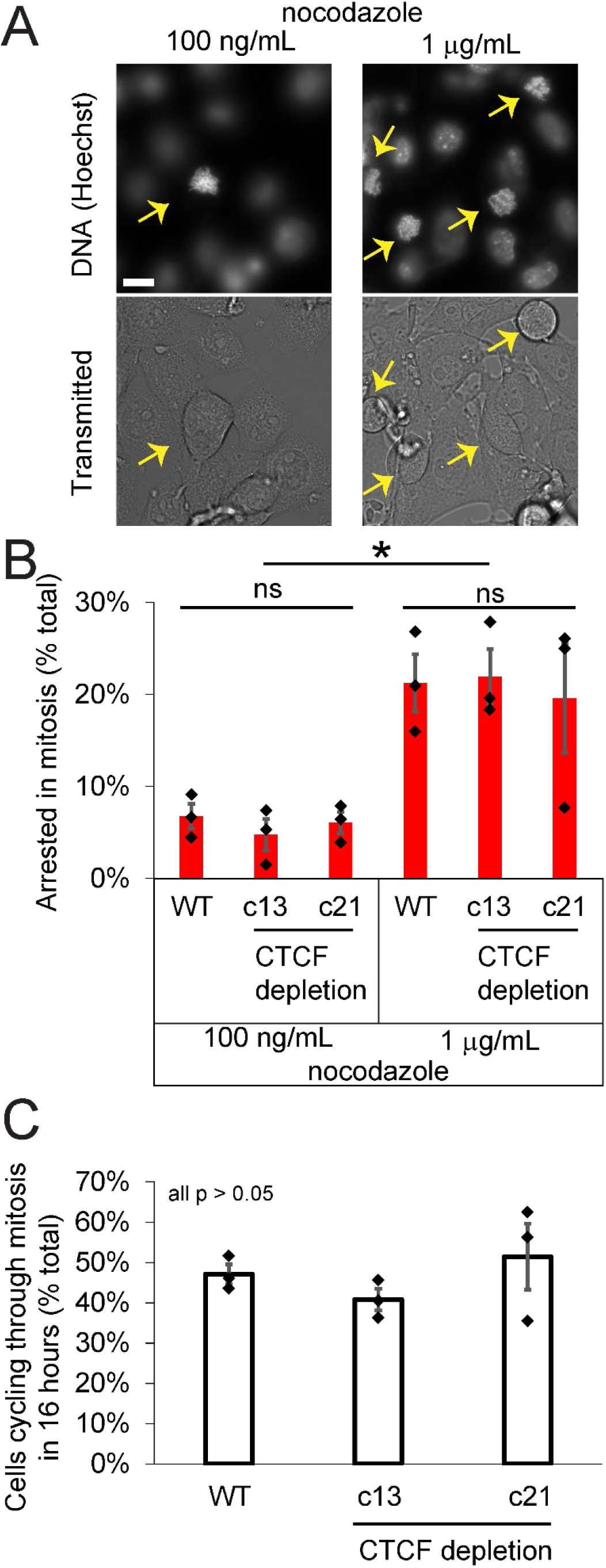
Cell cycling and activation of the mitotic checkpoint remains intact in CTCF knockdowns. (A) SiR-DNA labeled and brightfield transmitted light representative population images of B16 wild type (WT) and CTCF CRISPR knockdown clones (CTCF KD c13 and c21) treated with different concentrations of nocodazole. Yellow arrows denote cell in mitosis. (B) Graph of the percentage of cells in mitosis treated with 100 ng/mL or 1 µg/mL of nocodazole for 16 hours. (C) Graph of the percentage of untreated B16 wild type (WT) and CTCF CRISPR knockdown mutants (c13 and c21) cells that went through mitosis in 16 hours. Six technical replicates of n cells (WT 100 ng/mL 112, 120, 153, 81, 111, 89; WT 1 µg/mL n = 62, 50, 41, 67, 52, 44; c13 100 ng/mL 94, 108, 66, 78, 101, 78; c13 1 µg/mL 49, 43, 51, 38, 27, 48; c13 100 ng/mL 31, 51, 76, 21, 30, 14; c21 1 µg/mL 16, 46, 13, 16, 16, 20). Error bar represents standard error. Statistical tests are Student’s t-tests, significance denoted by * p < 0.05, ** p < 0.01, and *** p < 0.001, while no significance is denoted by ns. Scale bar is 10 µm.

Alternatively, CTCF knockdowns could be displaying mitotic failures due to faster cell cycling. To address the possibility of altered cell cycling, we measured the percentage of cells going through mitosis for 16 hours. Roughly 60% of the wild type B16-F1 cells population were visualized going through mitosis in 16 hours. The rate of cell cycling through mitosis was similar for both CTCF knockdowns (**Figure 2C**). Thus, it appears that abnormal mitoses in CTCF knockdowns are not due to spindle checkpoint failure or altered mitotic cycling.

### Mitosis is prolonged in instances of mitotic failure

If the mitotic checkpoint was activated to fix errors in mitosis this would likely prolong the length of mitosis. To determine if mitotic duration was altered in CTCF knockdowns, we measured the length of time from the beginning of mitosis to the end of segregation. In wild type cells the average mitotic duration was 25 ± 2 minutes (**Figure 3 and Supplemental Table 1**). Interestingly, successful mitoses in both CTCF knockdowns remained similar to wild type at 27 ± 5 minutes for c13 and 25 ± 2 minutes for c21. However, failed mitoses specifically showed a significantly prolonged mitosis increasing to 53 ± 19 minutes for c13 and 37 ± 9 minutes for c21 (**Figure 3, A and B**). Specifically, prolonged mitoses showed a delay in metaphase, suggesting that the mitotic checkpoint was delaying in order to correct chromosome alignment and/or attachments (**Figure 3A**). Thus, live cell imaging reveals problems occurring during mitosis that cause delay ultimately result in mitotic failures.

**Figure 3.**
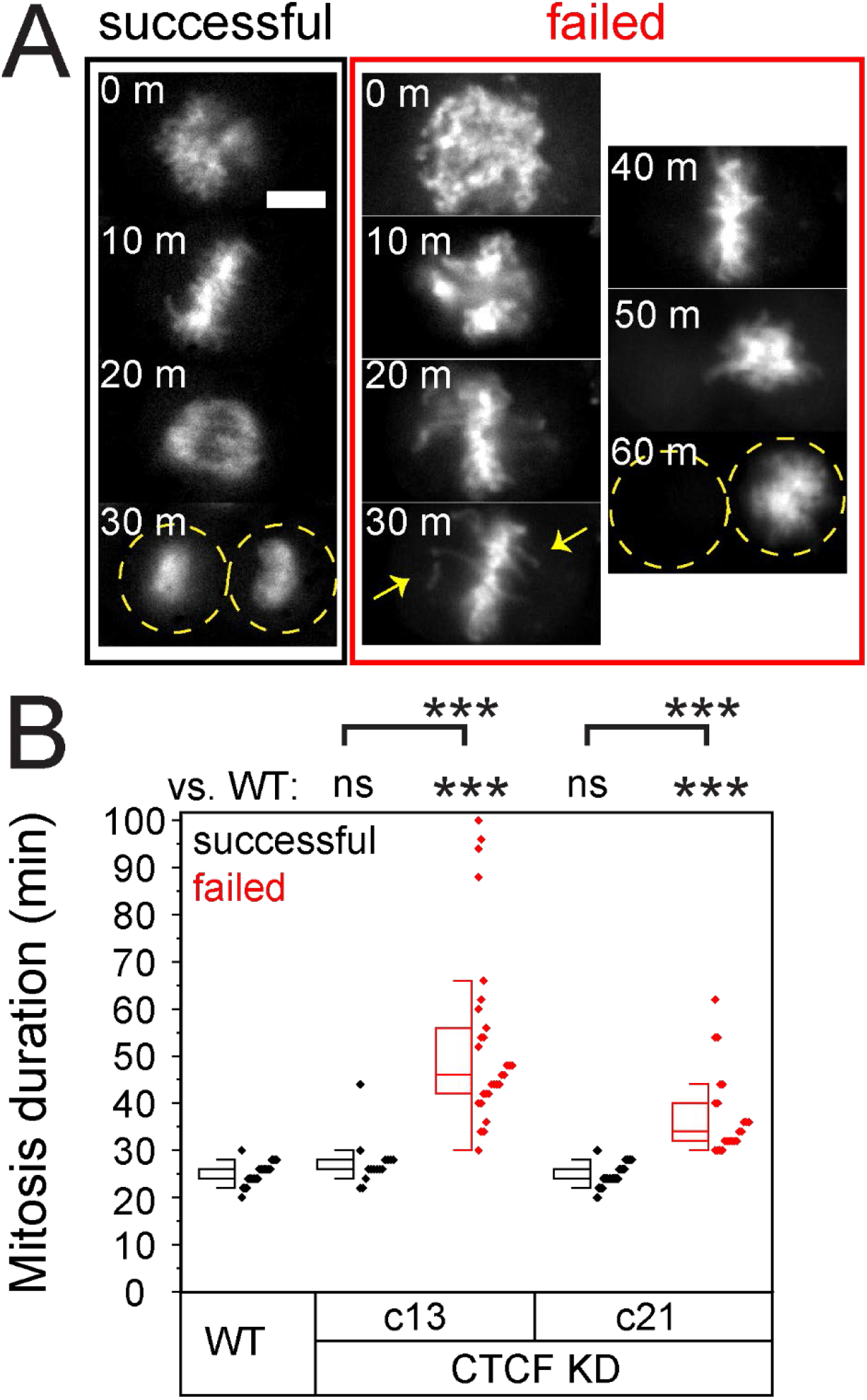
Failed mitoses display prolonged mitosis. (A) Example images of successful mitosis (black) and failed mitosis (red) labeled with SiR-DNA. Each image is 10 minutes later into mitosis, where most of the failed mitoses stalled in metaphase. Yellow arrows denote chromosomes of the metaphase plate. (B) Mitotic duration was measured for successful (black) and failed (red) mitoses and graphed as a box and whisker plot with individual events as small diamonds. Error bar represents standard error. Statistical tests are Student’s t-tests, significance denoted by * p < 0.05, ** p < 0.01, and *** p < 0.001, while no significance is denoted by ns. Scale bar is 10 µm.

### The mitotic spindle and chromosome organization are dependent on CTCF

Data showing prolonged mitosis in metaphase prompted us to investigate mitotic spindle structure. To determine if CTCF mutants cause mitotic failure through aberrant mitotic chromosome and spindle structure, we imaged DNA and microtubules via immunofluorescence. Consistent with time lapse data, we find that CTCF knockdowns result in aberrant spindle organization and structure. Wild type cells did not present abnormal nuclear spindles in all observed cases (n = 48, **Figure 4**). Oppositely, CTCF knockdowns displayed abnormal or and tri/tetrapolar spindles in 20-30% of all observed mitotic events (**Figure 4, A and B, and Supplemental Figure 1**). Abnormal mitotic spindle and chromosome organizations were defined largely by chromosomes organized behind the spindle pole. To quantify this abnormal behavior, we measured the edge of DNA relative to the spindle pole characterized by the brightest pixel of a-tubulin fluorescence. This was averaged over both poles (white arrow examples **Figure 4A** and graphed in **Figure 4C**). CTCF knockdown c13 mitotic spindles showed a significant change in DNA position relative to the spindle pole compared to wild type while c21 did not show significant change (**Figure 4C**). However, to determine if DNA was misplaced behind the spindle pole in individual mitotic spindles, we set a threshold of > 1 µm. With this threshold, we report that 0% of wild type spindles show DNA behind the spindle pole while 12% and 30% of CTCF knockdowns c13 and c21, respectively, show DNA behind the spindle pole (magenta diamond, **Figure 4C**). Therefore, CTCF is essential for the proper organization of the miotic spindle and chromosomes during mitosis.

**Figure 4.**
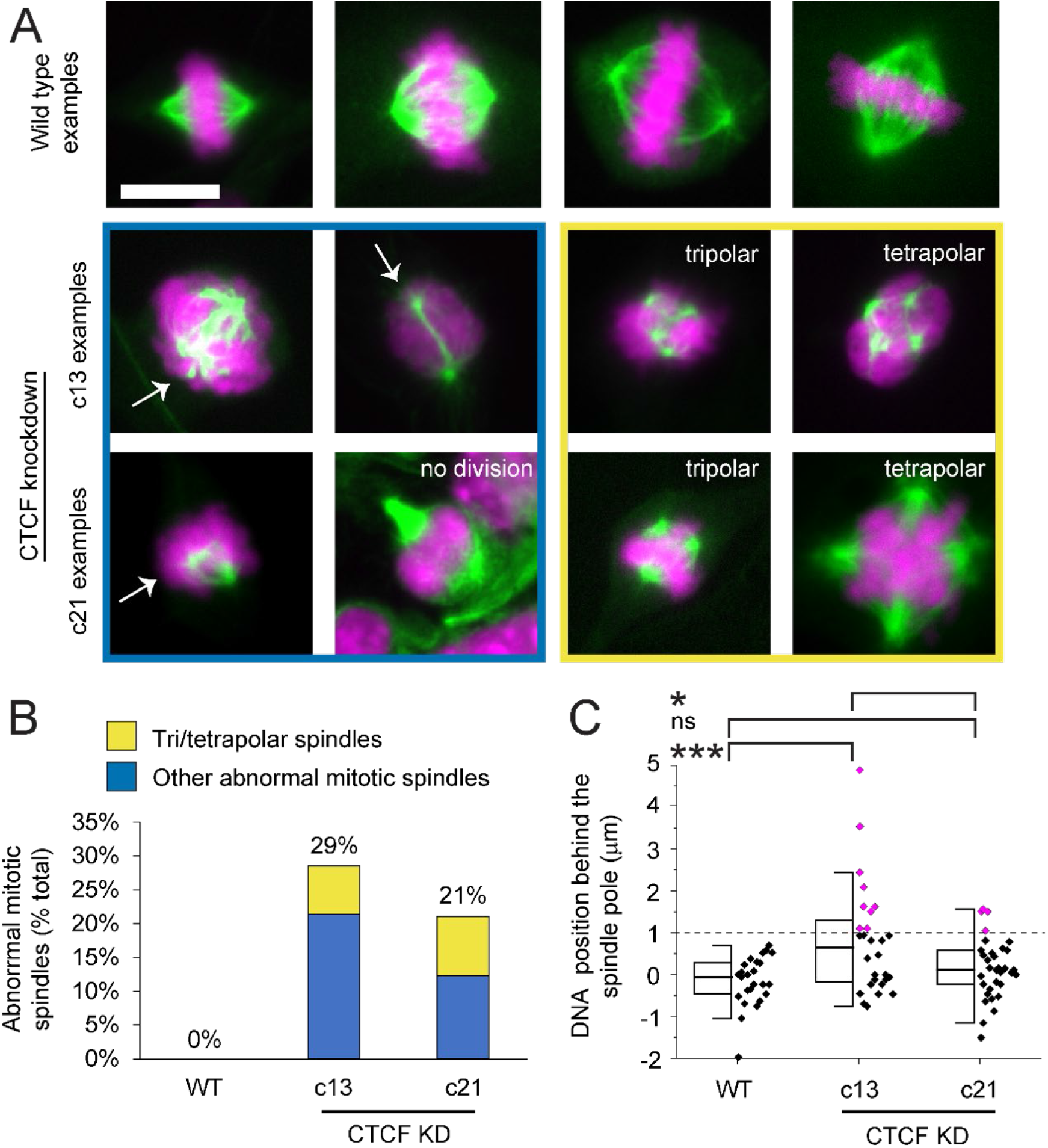
Mitotic spindle structure is perturbed in CTCF knockdowns. (A) Example images of mitosis in wild type and CTCF knockdowns c13 and c21 with labeled microtubules (green, α-tubulin) and DNA (magenta, Hoechst). Mitotic spindles were classified as abnormal mitotic spindle if they were tri-/tetrapolar spindles (yellow) or other abnormal (blue) if DNA was substantially behind the spindle pole or no division was observed. For all images see **Supplemental Figure 1**. (B) Graph of the percentage of abnormal mitotic spindles (WT n = 0/48; CTCF knockdown c13 n = 12/42; and CTCF knockdown c21 n = 12/57). (C) Box and whisker plot of average and individual measurement of DNA position relative to the spindle pole. Purple dots represent spindles where DNA is > 1 µm behind the spindle pole (WT 0%, c13 30%, c21 12%). Error bar represents standard error. Statistical tests are Student’s t-tests, significance denoted by * p < 0.05, ** p < 0.01, and *** p < 0.001, while no significance is denoted by ns. Scale bar is 10 µm.

### CTCF knockdowns present larger nuclei

Failures in mitosis can have lasting effects on interphase nuclei. We reasoned that our measured increase in anaphase segregation failures of the duplicated genome should lead to larger tetraploid nuclei. Thus, we measured nuclear size via area and DNA content via Hoechst sum intensity. As expected, we find that CTCF knockdowns result in an enrichment of large nuclei with high levels of DNA content compared to wild type cells (**Figure 5, A-C**). Average nuclear size nearly doubles from 150 ± 2 µm^2^ to 270 ± 10 µm^2^ in c13 and 265 ± 15 µm^2^ in c21 CTCF knockdowns (dashed lines, **Figure 5C**). Taken together, loss of CTCF results in anaphase segregation failure which results in larger nuclei with increased DNA content.

**Figure 5.**
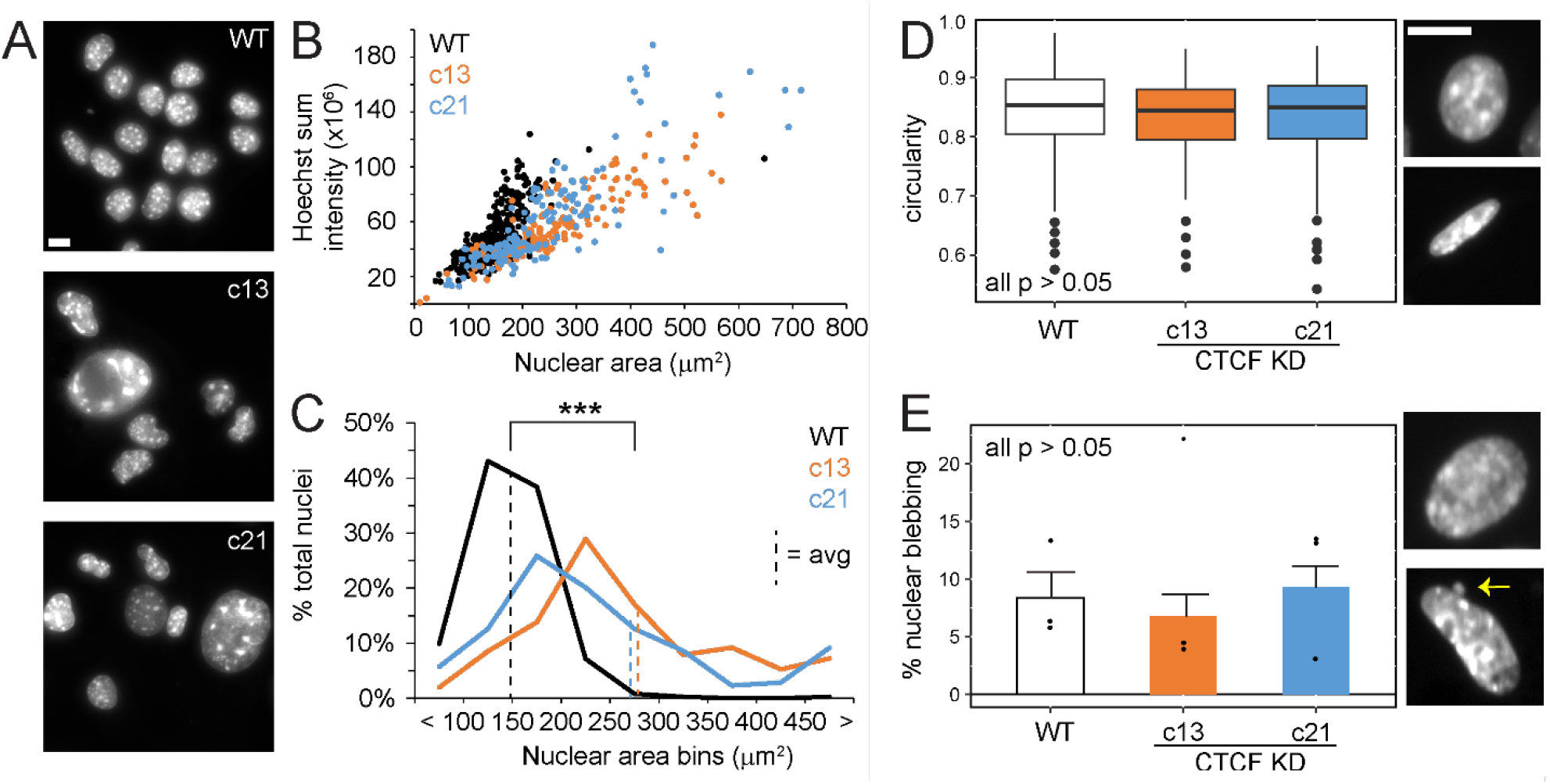
Nucleus size increases in CTCF knockdowns but circularity and nuclear blebbing do not change. (A) Example images of B16 nuclear sizes labeled with Hoechst. Graphs of (B) sum DNA intensity vs. nuclear area plotted for individual nuclei and (C) nuclear size distributions for wild type (black, n = 362) and CTCF knockdown clones c13 (orange, n = 151) and c21 (blue, n = 174). Dotted vertical line in panel C represents average nuclear size. (D)Graphs of nuclear circularity and (E) percentage blebbing for wild type (black, n = 35, 45, 64) and CTCF knockdown clones c13 (orange, n = 35, 27, 73) and c21 (blue, n = 35, 38, 65). (E) Graph of percentage blebbing for wild type (black, n = 65, 45, 64) and CTCF knockdown clones c13 (orange, n = 77, 27, 73) and c21 (blue, n = 144, 38, 65). Error bar represents standard error. statistical tests are Student’s t-tests, significance denoted by * p < 0.05, ** p < 0.01, and *** p < 0.001, while no significance is denoted by ns. Scale bar is 10 µm.

Changes in nuclear size and shape occur in many human diseases (Stephens et al. 2019a; Kalukula et al. 2022). Population measurements of nuclear shape via circularity and nuclear blebbing do not change between wild type and CTCF knockdowns (**Figure 5, D and E**). However, it should be noted that wild type B16-F1 nuclei already present a high incidence of decreased nuclear circularity 0.84 ± 0.01 and increased nuclear blebbing 8% compared to relative to other cell lines (Strom et al. 2021; Pho et al. 2022). Thus, wild type abnormal nuclear morphologies may hide underlying behaviors of mitotic failures in CTCF mutants. Overall, CTCF knockdowns do not cause abnormal nuclear morphology across the population.

### Anaphase failure in CTCF knockdowns directly results in abnormally shaped nuclei

Mitotic failure has been shown to cause disruptions to interphase nuclear morphologies (Gisselsson et al. 2001; Ohshima 2008). To directly determine if mitotic failures in CTCF knockdowns cause abnormal nuclear morphology, we tracked successful and failed mitoses via SiR-DNA and measured nuclear circularity in daughter nuclei 30 minutes post-mitosis (**Figure 6A**). First, we measured nuclear circularity in daughter nuclei of successful wild type mitoses, which revealed an average circularity of 0.95 ± 0.01. Interestingly, successful mitoses in both CTCF knockdowns also resulted in a similar circularity as wild type (0.93 ± 0.01, **Figure 6B**). However, in failed mitoses of CTCF knockdowns, the resulting daughter nuclei displayed a significantly decreased nuclear circularity to 0.77 ± 0.03 for c13 and 0.84 ± 0.02 for c21. This data suggests that post-successful-mitosis, nuclear shape is relatively circular while mitotic failures are associated with abnormal nuclear shape. This drastic outcome is likely hidden in population images where wild type B16-F1 nuclei undergo loss of nuclear shape throughout interphase as population measurements for wild type show a decreased circularity relative to 30 minutes post mitosis (0.84 ± 0.01 interphase population vs. 0.95 ± 0.01 successful mitosis daughter nuclei, p < 0.001). Thus, knockdown of CTCF results in mitotic failure that leads to abnormal nuclear morphology post-mitosis.

**Figure 6.**
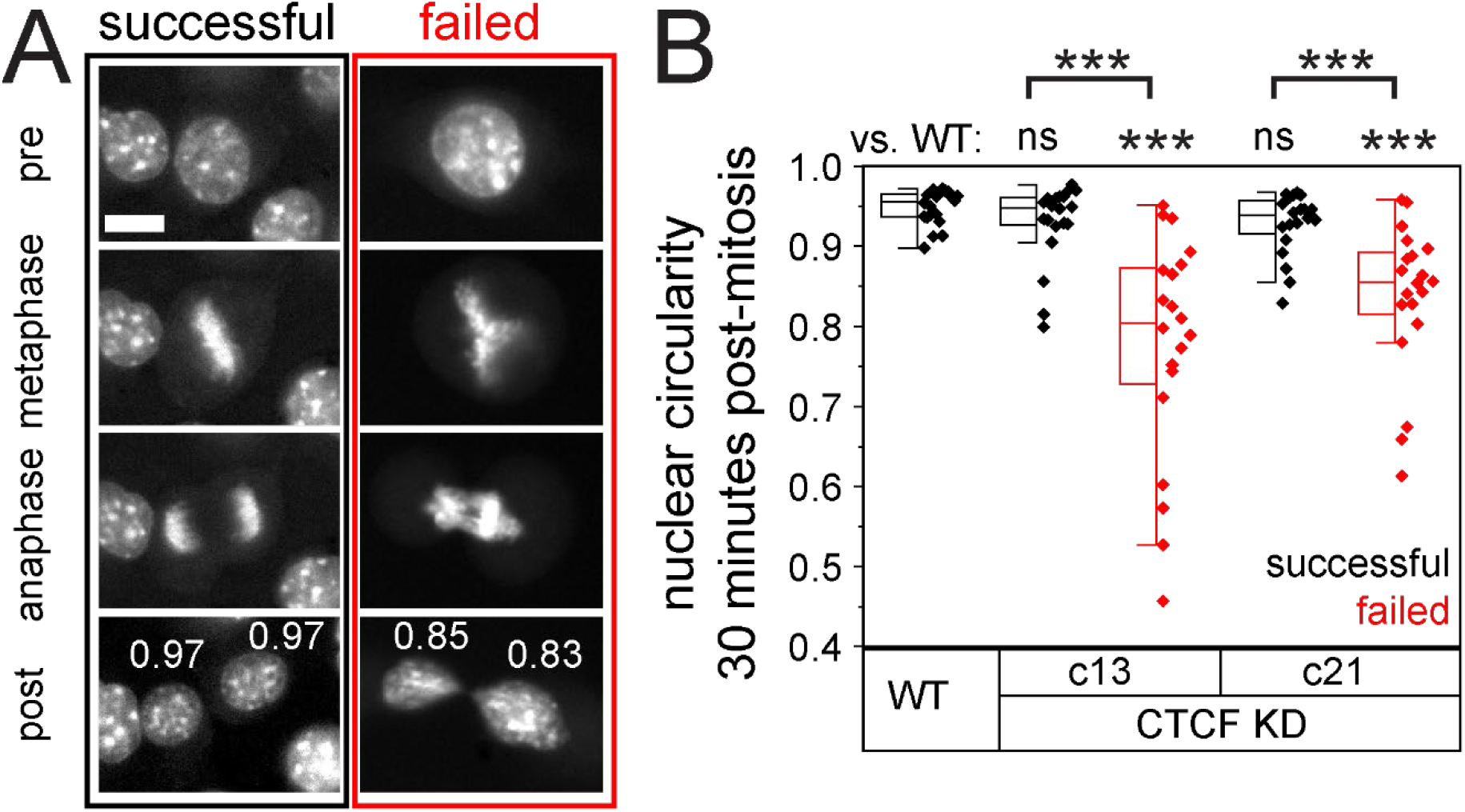
Mitotic failure causes abnormal nuclear shape post mitosis. (A) Example images of B16 successful (black) and failed (red) mitosis. Last image is 30 minutes post-mitosis where nuclear circularity was measured and noted in white. (B) Box and whisker plot and individual measurements of nuclear circularity 30 minutes post-mitosis for wild type (WT) and CTCF CRISPR knockdown mutants (c13 and c21) separated for successful (black) and failed (red) mitoses (n = 20 daughter nuclei per condition). Student’s t-tests, significance denoted by * p < 0.05, ** p < 0.01, and *** p < 0.001, while no significance is denoted by ns. Scale bar is 10 µm.

## Discussion

Here we detail CTCF is essential for proper mitosis in both metaphase spindle organization and anaphase segregation. This finding provides novel importance for CTCF beyond its highly studied role in chromatin genome organization via loops (Splinter et al. 2006; Hansen et al. 2017; Agbleke et al. 2020). Thus, while numerous studies have perturbed CTCF, few have considered the effects on mitosis and how that might impact chromatin and nuclear structure in interphase. Our data reveal that CTCF plays an essential part in mitotic fidelity which impacts the amount of chromatin in the interphase nucleus as well as size and shape.

### CTCF’s role in mitosis is supported by other studies

Past and present studies provide support for our finding that CTCF is essential for proper mitosis. A striking example is that mouse oocytes depleted of CTCF show failures in oocyte meiosis and in the embryo mitosis (Wan et al. 2008). Specifically, failures in mitosis resulted in larger nuclei, suggesting the genome did not segregate which recapitulates our findings in B16-F1 cells (**Figures 1 and 5**). In a CRSIPR gene screen of Hela cells CTCF knockouts showed tripolar spindles and increased DNA mean and integrated values suggesting increased amounts (Funk et al. 2022). This agrees with our findings of tri and tetrapolar spindles and increased DNA levels due to failed mitotic segregation in anaphase (**Figures 4 and 5**). Taken together, our and others novel findings support an essential role of CTCF in mitosis and a key revelation that CTCF is more than an interphase chromatin protein.

### The usual suspects for possible key CTCF interactors in mitosis

The mechanism for CTCF’s function in mitosis is likely complex. CTCF knockdowns result in disrupted spindle structure, chromosome organization, and anaphase segregation. This mix of varied effects suggests that CTCF has multiple roles in mitosis that could be due to its direct function and/or possibly through interactions with other major proteins in mitosis.

It has been reported that CTCF recruits CENP-E, a kinesin 7 kinetochore motor, to the pericentromere making it a possible mechanism for mitotic failure in CTCF knockdown cells (Xiao et al. 2015). Depletion of CENP-E (Putkey et al. 2002; Tanudji et al. 2004; She et al. 2020) or use of the CENP-E inhibitor GSK923295 (Bennett et al. 2015) causes errors in mitosis via effects mostly in metaphase spindle and chromosome disorganization. CENP-E disruption via CTCF knockdown might explain abnormal spindle structure, such as tri and tetra polar spindles, and chromosomes behind spindle poles and an intact spindle assembly checkpoint (She et al. 2020)(**Figures 2 - 4**). However, CENP-E perturbations clearly show individual polar chromosomes in metaphase spindles and clear missegregation of individual chromosomes during anaphase, which are seen infrequently in CTCF knockdowns. Furthermore, CENP-E does not present a cut phenotype of failed segregation during anaphase, a major phenotype of CTCF knockdowns (**Figure 1**). Alternatively, CTCF could have more roles than just recruiting CENP-E to the perceinteromere spring essential to mitosis (Gilbert et al. 2022). Therefore, CTCF disruption of CENP-E is a plausible mechanism but is poorly supported by the gross behaviors of CTCF knockdowns.

CAML-/- MEFs anaphase failure shows some of the same hallmarks of CTCF knockdown, but not all (Liu et al. 2009). In CAML-/ -cell mitosis presented a cut phenotype of failed anaphase segregations. Specifically, it is reported that CAML-/- cells, similar to our data for CTCF knockdowns, show spindle collapse upon anaphase onset leading to failed segregation and tetraploid cells. However, clear differences remain as CAML-/- cells were reported to have an effect on the spindle assembly checkpoint while not delaying metaphase, both opposite behaviors of CTCF knockdowns.

The obvious yet unclear possibility is that CTCFs has essential partnerships with other chromatin proteins or anaphase promoting proteins. Specifically, the cut phenotype, first detailed in yeast (Samejima et al. 1993), where non-segregating chromosome mass in anaphase is split by cytokinesis was reported to be due to failure of cohesin cleavage by separase (Hauf et al. 2001). It is possible that the loss of the N-terminal domain in these CTCF knockdowns (Kaczmarczyk et al. 2022) disrupts important interactions for both cohesion retention and chromatin loops (Pugacheva et al. 2020). The cut phenotype is one of the prominent displayed by both CTCF knockdowns. Other cut phenotype proteins of interest include topoisomerase 2 and condensing (Matsusaka et al. 1998). An alternative hypothesis is that CTCF aids the pericentromeric spring built by many chromatin proteins listed here to aid proper mitosis (Ribeiro et al. 2009; Stephens et al. 2011; Lawrimore and Bloom 2019). Thus, CTCF might support the function of these factors or directly contribute the larger functions of chromatin condensation and disentanglement.

### CTCF knockdown mitotic failures result in aberrant chromatin and nuclear interphase outcomes

CTCF’s role in mitosis is important because it causes direct effects on the interphase chromatin and nucleus. It has long been reported that mitotic failure can cause abnormal alterations to chromatin and nucleus in interphase (Gisselsson et al. 2001; Ohshima 2008; Flynn et al. 2021). The most straightforward of these is aneuploidy or changes in chromosome number. In CTCF knockdowns aneuploidy occurs due to the cut phenotype or an inability to segregate the genome which results in large changes in chromosome number. The most drastic forms of aneuploidy from complete segregation failure results in tetraploid cells which occurs frequently in CTCF knockdown cells (**Figure 1 and 5**). Having four copies of the genome causes massive changes in chromatin and transcriptional behavior and is a common disease state in cancer (Tanaka et al. 2018). Beyond this, tetraploid cells result in larger nuclei. It is also possible that tetraploid cells could have an abnormal nuclear mechanics via chromatin’s contribution (Stephens et al. 2017; Banigan et al. 2017; Hobson et al. 2020; Currey et al. 2022) that would disrupt many other important mechanisms in nuclear mechanobiology including mechanotransduction (Kalukula et al. 2022).

Mitotic failures result in abnormally shaped nuclei, a common cause and hallmark of human disease (Gisselsson et al. 2001; Stephens et al. 2019a). Abnormal nuclear morphology is well-documented to result in nuclear ruptures due to the high curvature of nuclear shape (Denais et al. 2016; Raab et al. 2016; Stephens et al. 2018; Xia et al. 2018; Strom et al. 2021; Pfeifer et al. 2022). Nuclear ruptures are known to disrupt key nuclear functions of transcription (Helfand et al. 2012; Le Berre et al. 2012), DNA damage/repair (Furusawa et al. 2015; Denais et al. 2016; Raab et al. 2016; Xia et al. 2018; Stephens et al. 2019b), and cell cycle control (Pfeifer et al. 2018). Recently, we reported that changes in transcription affect nuclear ruptures (Pho et al. 2022) where CTCF knockdown failures resulting in tetraploid nuclei may have highly altered transcription profiles. Thus, CTCF-based miotic failures can lead to nuclear dysfunction outside of CTCFs highly studied role in interphase chromatin looping.

### Conclusion

Even though we have studied mitosis for decades, we continue to find new contributors. CTCF is a well-known interphase chromatin protein that has been at the center of one of the greatest revolutions in our understanding of genome organization and function via long range looping. However, even though countless studies have investigated its interphase function, we detail that CTCF is essential to mitotic function and resulting interphase nucleus and chromatin properties. Further studies are needed to identify the role of CTCF in mitosis where it uniquely affects spindle structure, metaphase alignment, and anaphase segregation.

## Materials and Methods

### Cell Culture

B16-F1 cells, wild type (WT) and CTCF knockdown mutants (c13 and c21) were cultured in DMEM (Corning) completed with 10% fetal bovine serum (FBS, Hyclone) and 1% penicillin/ streptomycin (Corning). Cells were grown and incubated in 37°C and 5% CO_2_, passaged every 2 to 3 days and kept for no more than 30 passages.

### Time Lapse Imaging

Time lapse images were acquired with Nikon Elements software on a Nikon Instruments Ti2-E microscope with Crest V3 Spinning Disk Confocal, Orca Fusion Gen III camera, Lumencor Aura III light engine, TMC CLeanBench air table, with 40x air objective (N.A 0.75, W.D. 0.66, MRH00401). Time lapse imaging was possible over a long duration using Nikon Perfect Focus System and Okolab heat, humidity, and CO2 stage top incubator (H301). Images were captured via 12-bit sensitive camera. Cells were treated with SiR-DNA (Cytoskeleton Inc.), plated and imaged in 4-well cover glass dishes (Cellvis). Images were taken in 2-minute intervals for 20 hours with 6 fields of view per experimental condition, using Cy5 and transmitted imaging modalities to capture DNA (via SiR-DNA), nuclear structure, and cell body structure.

### Immunofluorescence

Three cell lines of B16-F1 mouse melanoma cells were used: wild type cells (WT), cells with a CRISPR C13 deletion mutation in the CTCF gene (c13), and cells with a CRISPR C21 deletion mutation in the CTCF gene (c21). All live cells coverslips were fixed in 3.2% paraformaldehyde, 0.1% glutaraldehyde, and permeabilized with 0.5% Triton X-100 for 10 minutes. The coverslips were rinsed in PBS-TW-Az (phosphate buffered saline with 0.1% tween-20 and 0.02% sodium azide). Afterwards, humidity chambers were made using filter paper, distilled water, and parafilm. Coverslips were placed on 50 µL drops of the primary antibody mixture consisting of rat anti-alpha tubulin at 1:100 dilution (Novus biological); mouse anti-lamin A/C at 1:100 dilution (Cell Signaling Technology); and PBS with 5% BSA. Primary antibodies were incubated in a 37º C for 60 min. After incubation, each coverslip was washed 3 times in PBS then placed on top of 50 µL drops of the secondary antibody mixture of goat anti-rat TRITC (1:100; Millipore Sigma) and goat anti-mouse FITC (1:100; Millipore Sigma) in newly prepared humidity chambers. These dishes were again incubated at 37° C for 30 minutes. All coverslips were then washed 3 times in PBS before being mounted on a drop of mounting medium (DAPI fluoromount – G 1:1000; SouthernBiotech)) on labeled glass coverslips. To prevent cells from shearing, side-to-side movement was limited as much as possible. All coverslips were sealed with a small amount of Sally Hansen instant dry clear nail polish to be dried overnight.

### Immunofluorescence Imaging

A Nikon TE2000 inverted widefield fluorescence microscope was used to image cell lines with a Plan Fluor 40x objective. Images were taken on a QImaging Fast 1394 CCD camera using MicroManager, saved as Tif files, and later stacked with color on FIJI-ImageJ. Cells were first imaged using transmitted light to focus on the field of view and visualize overall cell morphology. Blue fluorescent light (excitation 480 nm) was used to visualize lamin AC, green fluorescent light (excitation 560 nm) was used to visualize alpha-tubulin, and UV light (excitation 360 nm) was used to visualize DAPI stained nuclei. Wild-type (WT) slides were observed first to understand standard mitotic cells to later compare to the abnormal mitotic structures in CRISPR-depleted clone cells (c13 and c21). In FIJI-ImageJ, green was applied to the alpha-tubulin, and magenta was applied to the chromatin.

### Mitotic spindle and chromosome organization analysis

Images of mitotic spindles were analyzed using FIJI-ImageJ. Images were categorized as normal, tri/tetrapolar spindle, or abnormal spindle, based on the appearance of DNA outside the edge of the spindle. Cells exhibiting non-tri/tetrapolar spindle abnormalities along the metaphase plate were compiled and the average distance of DNA that resided beyond the spindle poles was quantified via line scan analysis in FIJI ImageJ. Line scans were drawn from each spindle pole to an arbitrary point beyond the final edge of the cell. Based on the Plot Profiles of the line scans, the peaks of the green and magenta channels were identified, and half maximum calculated, accounting for Gaussian blur of spindle poles (full-width-half-maximum). Their corresponding pixel values were used as the locations of the spindle and DNA. The distance between the DNA and spindle was calculated by finding the difference between these two-pixel values.

### Nuclear Population Analysis

Images were analyzed using Fiji, set scale measurement distance in pixel was 8.6 pixel/µm by known distance 1, pixel aspect ratio 1, unit of length 1, then selected Global. For each field of view, each cell’s parameter was circled to produce an ROI of each cell. From ROI data, nuclear size was collected using ROI area size measurement in units of µm^2^. DNA content was measured using Hoechst intensity in arbitrary units of hundreds of thousands. Measurements for solidity and circularity for each cell’s nucleus were also gathered from ROI information. Cells that appeared visually abnormal as well as normal were compiled into one data set. Blebs were counted for each field of view. Blebs were counted for each field of view with a threshold value of >1 µm in size.

### Mitotic Time Lapse Events Analysis

Time lapse images were saved within the NIS-Elements AR software for observation and analysis. Images were observed to record each individual mitotic event that transpired over the time lapse. Whether a mitotic event was successful or failed was recorded, as well as the type of mitotic failure that was observed during a failed mitotic event. Once this data was collected across WT and the two mutants, it was compiled and averaged for each condition to provide further analysis in Excel (Microsoft) to determine % failed mitotic events and the breakdown percentages for each type of mitotic failure recorded for each condition. Statistical significance was determined via *t* tests between the conditions.

Time lapse data was further analyzed to determine duration of mitotic events in minutes for both successful and failed. To achieve this, initial frame of mitosis, initial frame of anaphase, and frame of cytokinesis were recorded in Excel (Microsoft) for each mitotic event. Initial frame of mitosis was determined as the frame when nucleus began to lift and ball up, and frame of cytokinesis was determined visually through use of both SiR-DNA and brightfield. Once this data was collected, duration of mitosis was calculated by taking the difference between the cytokinetic frame and the initial frame of mitosis and multiplying it the length of time between frames which was 2 minutes. This data was then sorted between successful and failed mitotic events, averaged, and graphed in Origin. Statistical significance was determined through use of the *t* test.

Images were observed for mitotic events to record and analyze post-mitotic interphase shape across each condition. ROIs were hand-drawn around daughter nuclei 15 frames, or 30 minutes (2-minute interval between frames), post-mitosis using the Bezier ROI drawing tool. For the two CTCF depleted mutants, this procedure was done for successful and failed mitotic events. Due to the low percentage of failed mitotic events in WT, only daughters from successful mitotic events were measured. ROIs were converted to binary to extract circularity object data, which was then exported and compiled to Excel (Microsoft). Compiled data was then averaged and statistical significance between the conditions was determined via Student’s *t* tests.

## Acknowledgements

We would like to thank Patricia Wadsworth and Thomas Marasca for helpful and insightful discussions and general support. We would also like to thank HHMI which purchased microscopes used in Bioimaging class via a grant and The Biology Department at UMass Amherst for use of the facilities associated with ISB 360. KC, YB, NE, and ADS are supported by the Pathway to Independence Award (R00GM123195) and Center for 3D Structure and Physics of the Genome 4DN2 grant (1UM1HG011536). GG would like to thank for the Israel Cancer Association (grant no. 202220036) and Ariel University for their support. The authors declare no competing interests.

---

*Movie 1. Normal mitosis*.

*Movie 2. Anaphase fail segregation*.

*Movie 3. Anaphase fail no segregation*.

*Movie 4. Metaphase fail no segregation*.

*Movie 5. Tripolar segregation*.

*Supplemental Table 1. Raw data*. Excel document with the raw data for each figure.

*Supplemental Figure 1. Individual images of mitotic spindles*. The individual images used to score Figure 4. Yellow outlines denote abnormal metaphase spindles besides tri or tetra polar spindles. DNA vs. spindle pole relative distance measures can be found in Supplemental Table 1.

