## Supplemental Figure 1 for "CTCF is essential for proper mitotic spindle structure and anaphase segregation"

### Combined WT (PMAT)

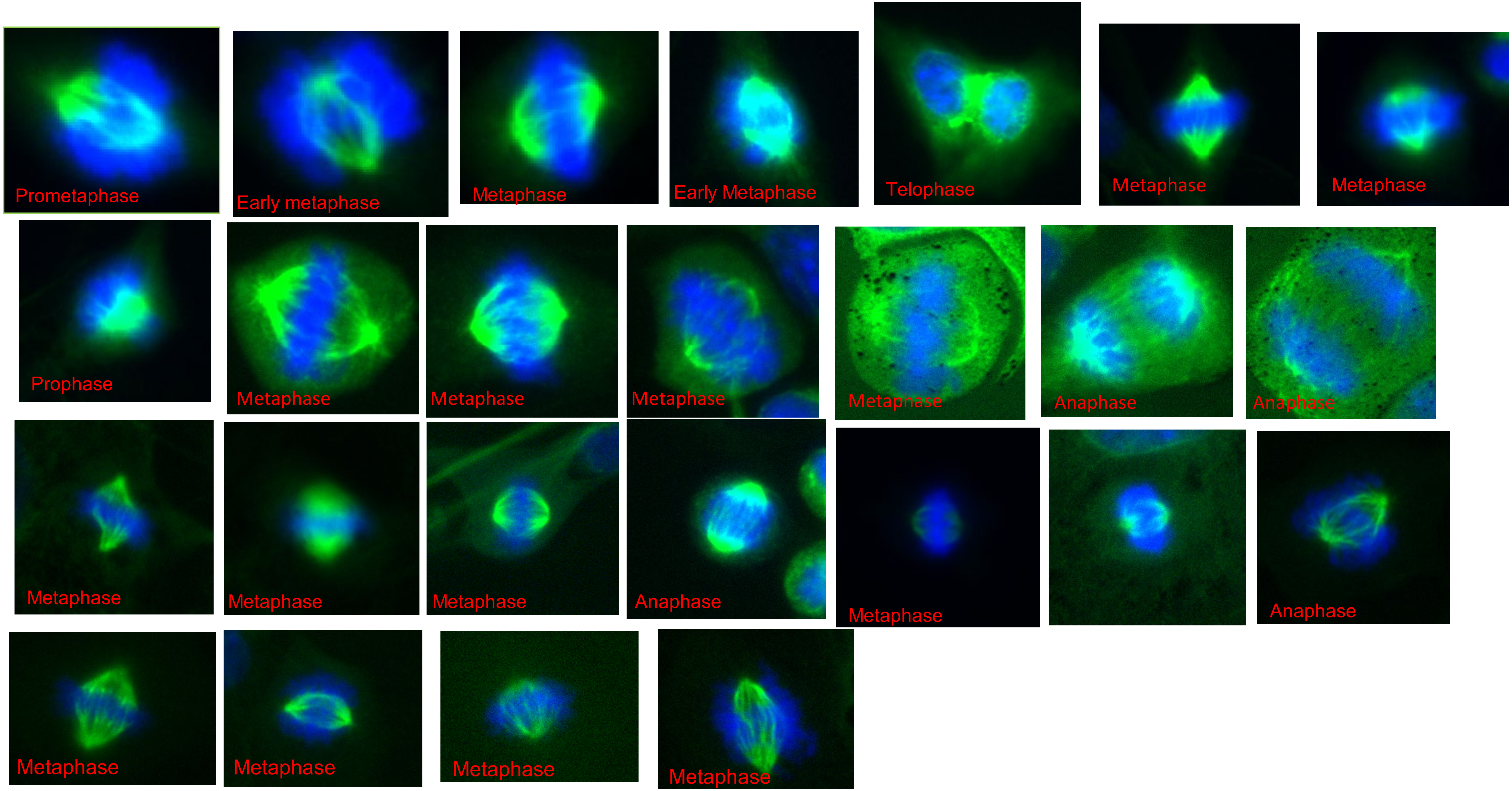

### Combined WT (cytokinesis)

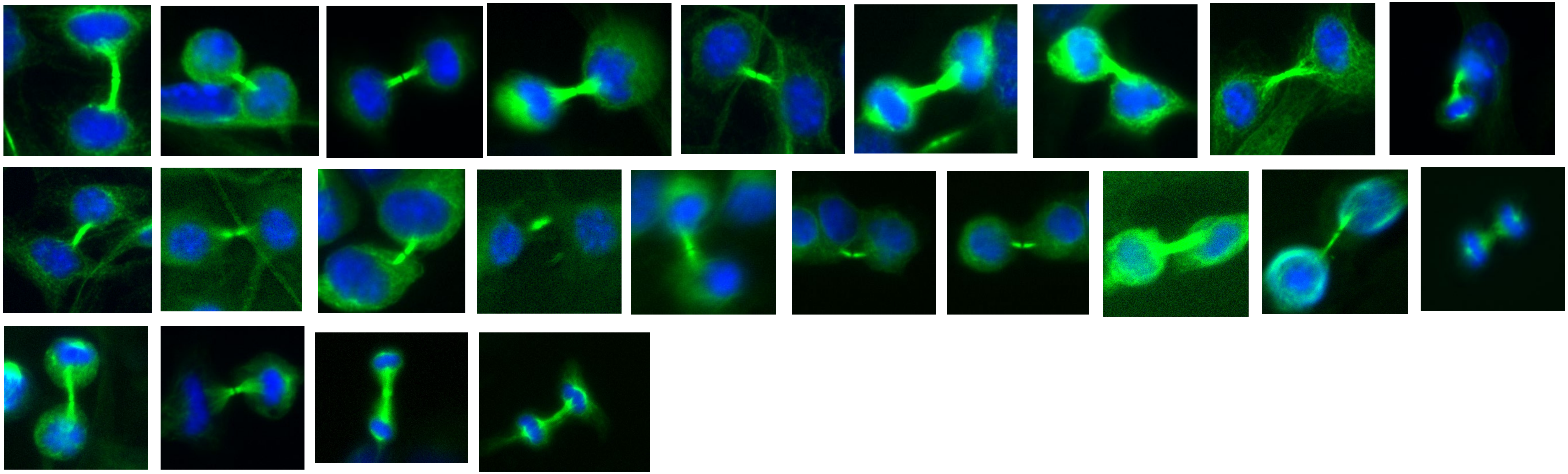

### Combined CTCF KD c13

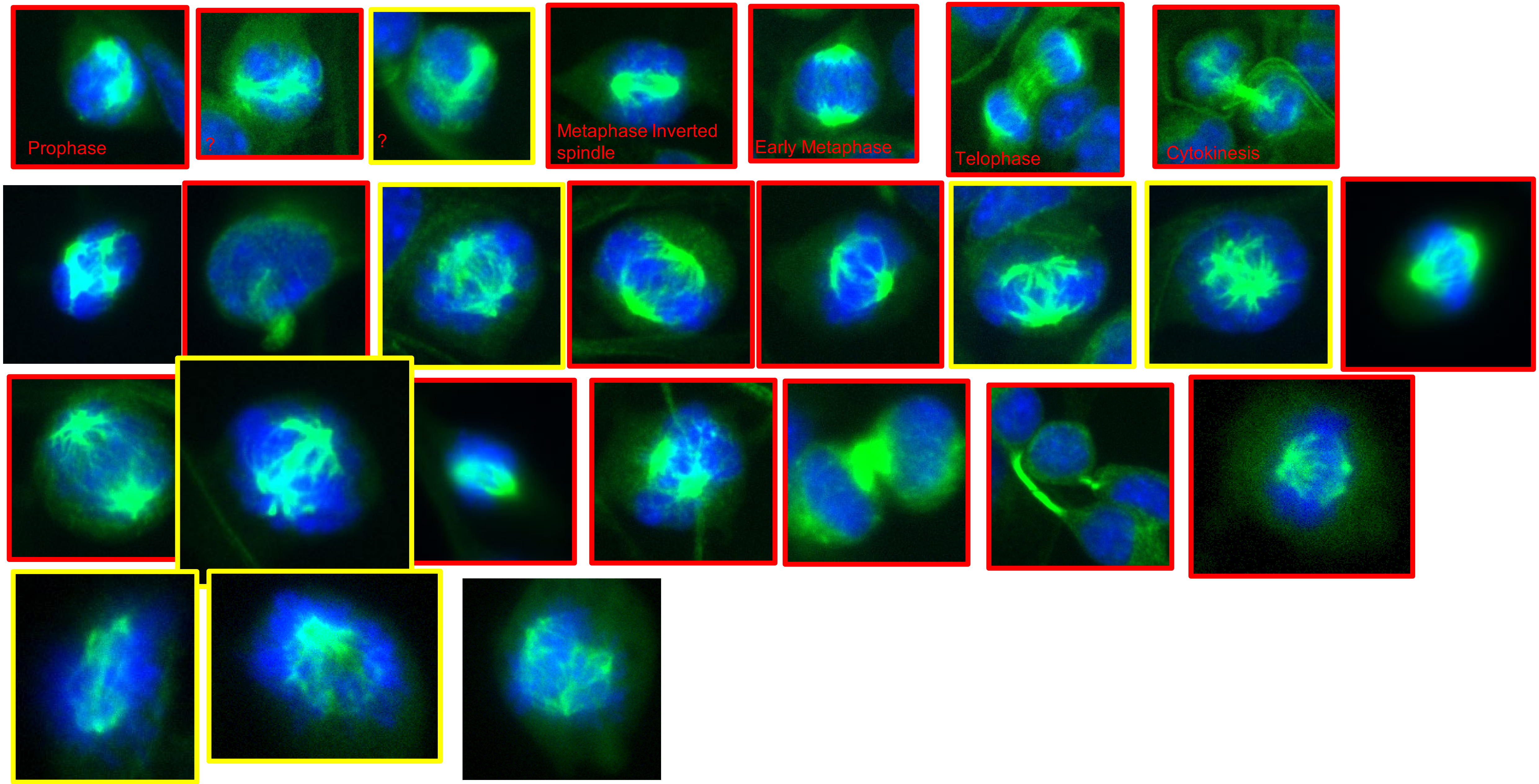

### Combined CTCF KD c13 cont.

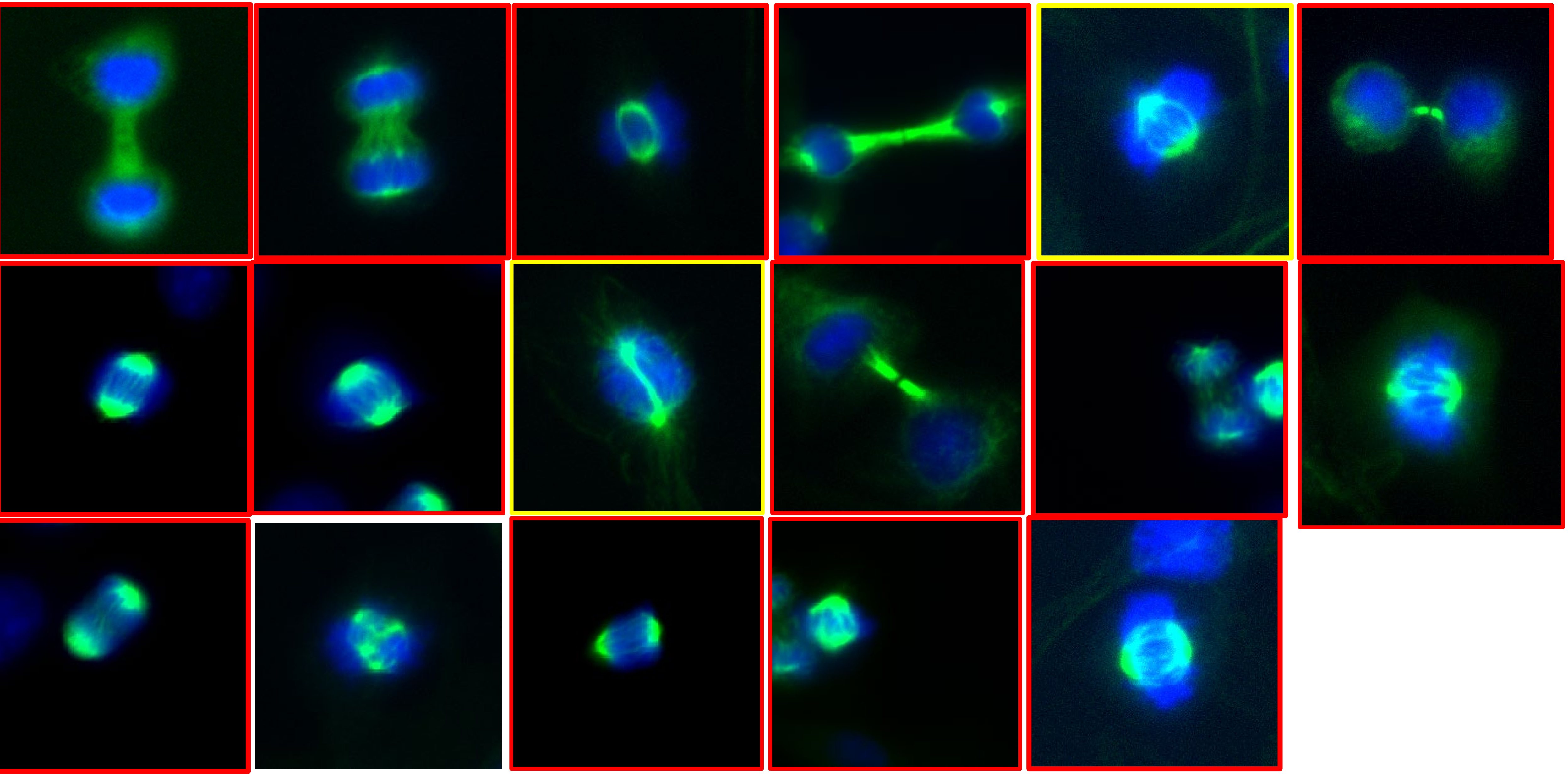

### Combined CTCF KD c21 (PMAT+C)

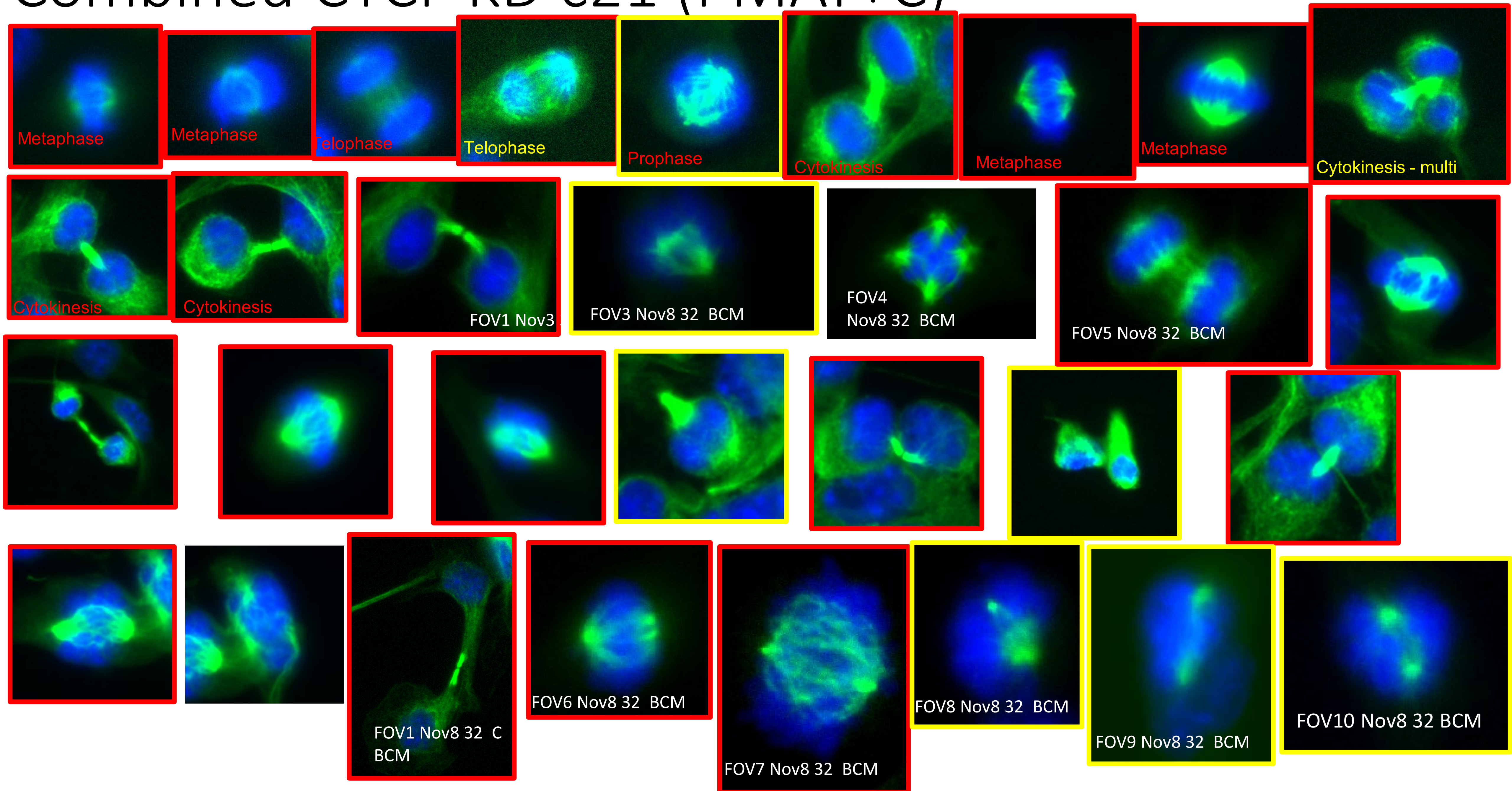

### Combined CTCF KD c21 (PMAT+C) cont.

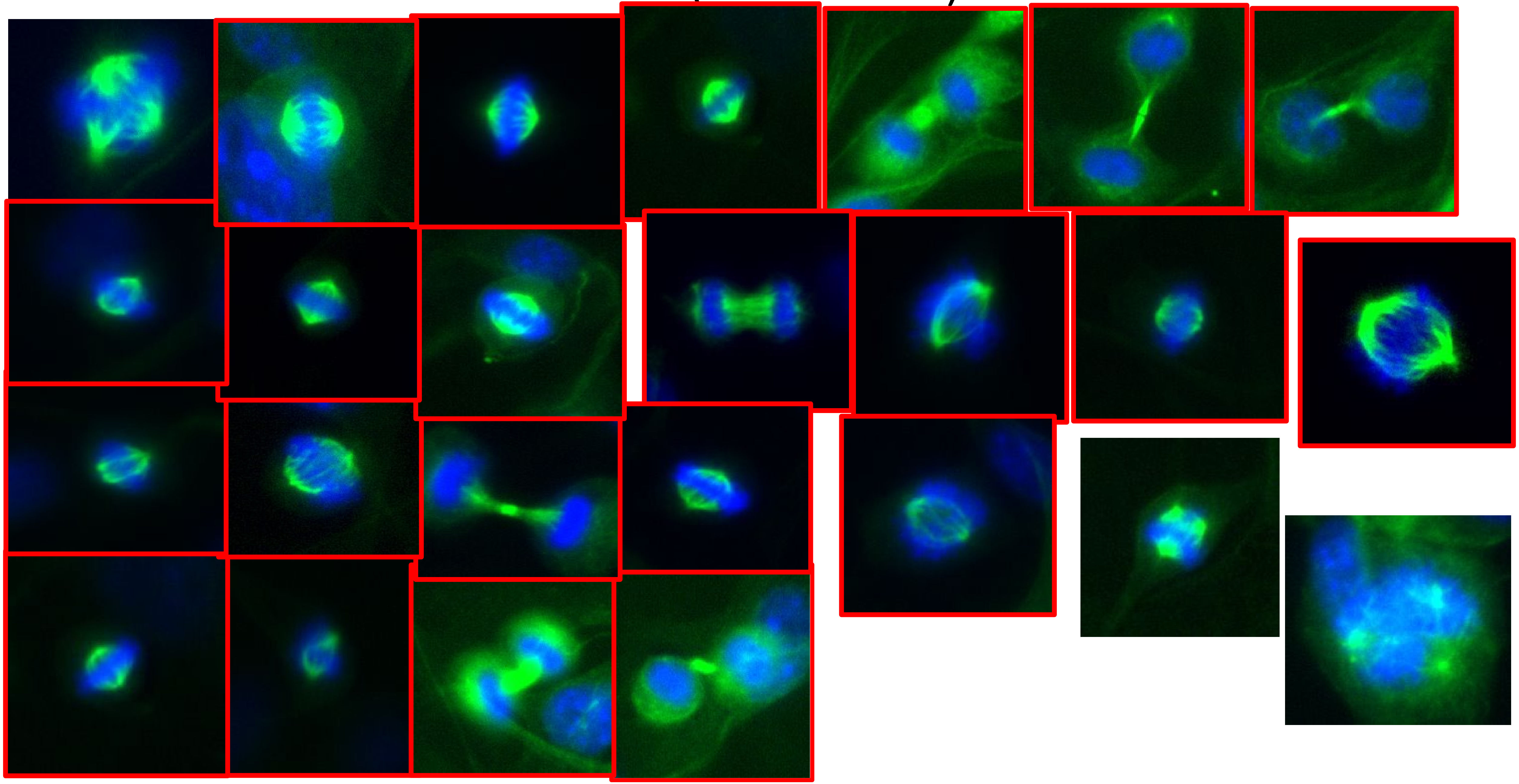
